# Size-dependent patterns of cell proliferation and migration in freely-expanding epithelia

**DOI:** 10.1101/2020.02.28.970418

**Authors:** Matthew A. Heinrich, Ricard Alert, Julienne M. LaChance, Tom J. Zajdel, Andrej Košmrlj, Daniel J. Cohen

**Affiliations:** Department of Mechanical and Aerospace Engineering, Princeton University, Princeton, NJ 08544, USA; Lewis-Sigler Institute for Integrative Genomics, Princeton University, Princeton, New Jersey 08544, USA; Princeton Center for Theoretical Science, Princeton University, Princeton, New Jersey 08544, USA; Princeton Institute for the Science and Technology of Materials (PRISM), Princeton University, Princeton, New Jersey 08544, USA

**Keywords:** tissue expansion, cell cycle, collective cell migration, epithelia, active polar fluids

## Abstract

The coordination of cell proliferation and migration in growing tissues is crucial in development and regeneration but remains poorly understood. Here, we find that, while expanding with an edge speed independent of initial conditions, millimeter-scale epithelial monolayers exhibit internal patterns of proliferation and migration that depend not on the current but on the initial tissue size, indicating memory effects. Specifically, the core of large tissues becomes very dense, almost quiescent, and ceases cell-cycle progression. In contrast, initially-smaller tissues develop a local minimum of cell density and a tissue-spanning vortex. To explain vortex formation, we propose an active polar fluid model with a feedback between cell polarization and tissue flow. Taken together, our findings suggest that expanding epithelia decouple their internal and edge regions, which enables robust expansion dynamics despite the presence of size- and history-dependent patterns in the tissue interior.

## Introduction

Writing in 1859, physiologist Rudolf Virchow presented the concept of the ‘Zellenstaat’ or ‘Cell State,’ describing tissues as “a society of cells, a tiny well-ordered state” (1). This social framework motivated Abercrombie and Heaysman’s 1954 work on cellular behavior that elucidated how encounters between cells can regulate locomotion and proliferation via contact inhibition (2). Since then, concerted interdisciplinary effort has been brought to bear on understanding how cell-cell interactions give rise to the complex collective behaviors driving so many crucial biological processes. One of the most foundational collective behaviors is collective cell migration— the directed, coordinated motion of cellular ensembles that enables phenomena such as gastrulation, wound healing, and tumor invasion (3). Given this importance, considerable effort spanning biology, engineering, and physics has been directed towards understanding how local cellular interactions can give rise to globally coordinated motions (4, 5).

Studies of collective cell migration are most often performed using epithelial tissues due to their fundamental role in multicellular organisms and strong cell-cell adhesion, which in turn gives rise to elegant, cohesive motion. Moreover, given that epithelia naturally form surfaces in vivo, studying epithelial layers in vitro has a physiological basis that can inform our understanding of processes such as healing (6), envelopment (7), and boundary formation (8). These features have made epithelia both the gold standard in collective cell migration studies, and one of the most well-studied models for biological collective behaviors.

Due to the complexity of collective behaviors, much effort has gone towards reductionist assays that restrict degrees of freedom and ensemble size to simplify analysis and interpretation. One such approach is to confine a tissue within predefined boundaries using micropatterning to create adhesive and non-adhesive regions (9–14). Such confinement mimics certain in vivo contexts such as constrained tumors as well as aspects of compartmentalization during morphogenesis (15). Alternately, many studies have explored the expansion of tissues that initially grow into confluence within confinement but are later allowed to migrate into free space upon removal of a barrier. A popular assay of this type relies on rectangular strips of tissue that are allowed to expand in one or both directions (6, 16–23), where averaging along the length of the strip can reveal coordinated population-level behaviors such as complex migration patterns, non-uniform traction force fields, and traveling mechanical waves. Other studies have focused on the isotropic expansion of micro-scale (< 500 *μm* diameter) circular tissues using the barrier stencil technique (24) as well as photoswitchable substrates (25). Still more work has explored approaches to induce directional migration, from geometric cues to applied electric fields (26, 27).

In contrast to micro-scale confinement assays, other work has focused on large, freely-expanding tissues of uncontrolled initial size and shape, which grow from either single cells (28, 29) or cell-containing droplets (30, 31). Related experiments track long-term growth of cell colonies via images taken once per day over several days, but this low temporal resolution cannot access timescales over which migration is important (29, 32). Thus, there is still a lack of assays to study long-term expansion and growth of large-scale tissues with precisely-controlled initial conditions, especially initial tissue size, shape, and density.

To address this gap, we leveraged bench-top tissue patterning (6, 33) to precisely pattern macro-scale circular epithelia of two sizes (>1 mm in diameter) and performed longterm, high frequency, time-lapse imaging after release of a barrier. To elucidate the consequences of size effects on the tissue, we tracked every cell, relating the overall expansion kinetics to cell migration speed, cell density, and cell-cycle dynamics. We find that, whereas the tissue edge dynamics is independent of the initial conditions, the tissue bulk exhibits size-dependent patterns of cell proliferation and migration, including large-scale vortices accompanied by dynamic density profiles. Together, these data comprise the first comprehensive study of macro-scale, long-term epithelial expansion, and our findings demonstrate the importance of exploring collective cell migration across a wider range of contexts, scales, and constraints.

## Results

### Expansion of millimeter-scale epithelia of different sizes and shapes

We began by characterizing the overall expansion and growth of tissues with the same cell density but different initial diameters of 1.7 mm and 3.4 mm (a 4X difference in area, with tissues hereafter referred to as either “small” or “large”), using an MDCK cell line stably expressing the 2-color FUCCI cell-cycle marker (22, 31, 34–36). We patterned the tissues by culturing cells in small and large circular silicone stencils for ~18 hrs (6, 33), whereupon stencils were removed and tissues were allowed to freely expand for 46 h (Fig. 1A, Movie S1), while images were collected at 20 minute intervals using automated microscopy (see SI Supplementary Methods). Our cell seeding conditions and incubation period were deliberately tuned to ensure that the stencils did not induce contact inhibition or jamming prior to stencil removal. Upon stencil removal, tissues expanded while maintaining their overall circular shape throughout the 2-day experiment. Unless otherwise noted, cell density at stencil removal was ~2700 cells/mm^2^, a value consistent with active and growing confluent MDCK epithelia (22, 35). First, we measured relative areal increase (Fig. 1B) and relative cell number increase (SI Appendix, Fig. S1) of small and large tissues. By 46 h, small and large tissues had increased in area by 6.4X and 3.3X, respectively, while cell number increased by 9.2X and 5.5X, respectively (Fig. S1). Since proliferation outpaces area expansion in long-term growth, average tissue density increased by the end of the experiment. The evolution of average tissue density was more complex, however, as small tissues experienced a density decrease from 4-12 h while large tissues exhibited a monotonic increase in cell density (Fig. 1C). Accordingly, at any given time after stencil removal, large tissues had a higher density than small tissues. Non-monotonic density evolution has been observed in thin epithelial strips (6) and likely arises from competition between migration and proliferation dynamics, which we discuss later.

**Fig. 1.**
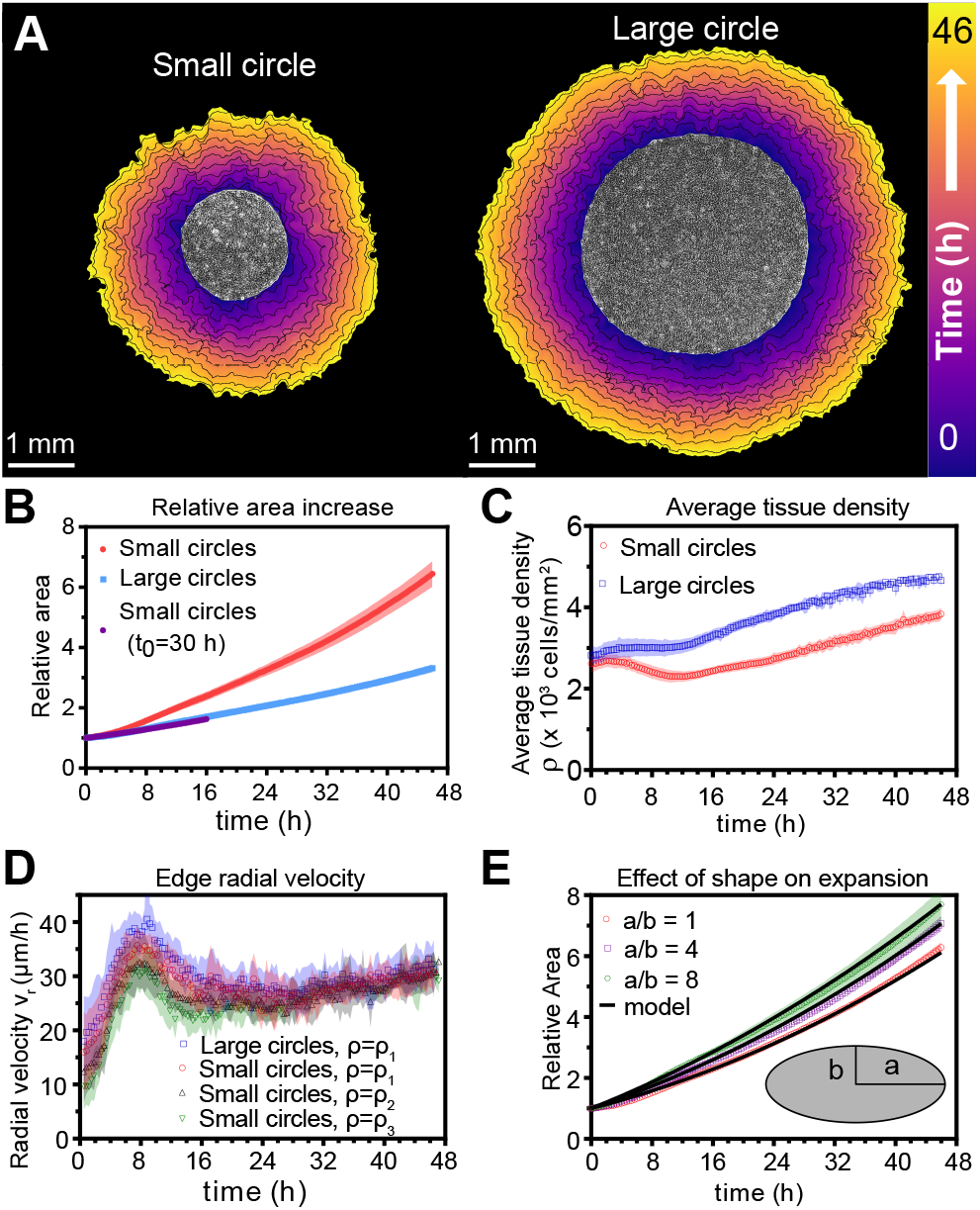
Expansion dynamics of millimeter-size cell monolayers. (A) Footprint throughout 46 h growth period of representative small (left) and large (right) circular tissues, with the tissue outlines drawn at 4 h increments. Initial diameters were 1.7 mm and 3.4 mm. (B) Small circles exhibit faster relative area, *A*(*t*)/*A*_0_, increase than large circles, where *A*_0_ and *A*(*t*) are the areas of tissues at the beginning of the experiment and at time *t*, respectively. Purple points show the relative area increase, *A*(*t* + *t*_0_)/*A*(*t*_0_), of small tissues from the time *t*_0_ = 30 h when they reached the size of the large circles. (C) Average tissue density *ρ*(*t*) = *N*(*t*)/*A*(*t*) has non-monotonic evolution in small tissues but monotonically increases in large tissues, where *N*(*t*) is the number of cells in a tissue at time *t*. (D) Edge radial velocity *υ_r_* is largely independent of initial tissue size and cell density. We grouped initial cell densities as *ρ*_1_ = [2350,3050] cells/mm^2^, *ρ*_2_ = [1650,2350) cells/mm^2^, and *ρ*_3_ = [1300,1650) cells/mm^2^. (E) Experimental data on tissue shape and model fits. Assuming a constant migration speed *υ_n_* in direction normal to the edge, we can predict the area expansion dynamics of elliptical tissues with different aspect ratios. Fitting the model to our data for all tissues gives *υ_n_* = 29.5 *μ*m/h. In B, data are from n=16 tissues across 5 independent experiments (small and large circles). In C, n=11 across 4 experiments for small circles), and n=9 across 3 experiments for large circles. In D, n=16 across 5 independent experiments for small and large circles, *ρ* = *ρ*_1_; n=13 across 3 experiments for small circles, *ρ* = *ρ*_2_; and n=11 across 3 experiments for small circles, *ρ* = *ρ*_3_. In E, n=4 across 2 experiments for a/b=1 and a/b=4, and n=5 across 2 experiments for a/b=8. Shaded regions correspond to standard deviations.

We then related area expansion to the kinematics of the tissue edge. To quantify edge motion, we calculated the average radial velocity of the tissue boundary, *υ_r_*(*t*), at 1 hr intervals over 46 hrs (SI Supplementary Methods). We found that *υ_r_* is independent of both tissue size and a wide range of initial cell densities, in all cases reaching ~30 *μ*m/h after ~16 h (Fig. 1D). Before reaching this constant edge velocity, *υ_r_* ramps up during the first 8 h after stencil removal, and, notably, overshoots its long-time value by almost 30%. We hypothesize that the overshoot is due to the formation of fast multicellular finger-like protrusions that emerge at the tissue edge in the early stages of expansion and then diminish (Movie S2). This hypothesis is supported by a recent model showing that edge acceleration (as observed during the first 8 h in Fig. 1D) leads to finger formation (37). It is remarkable that the edge radial velocity *υ_r_*(*t*) is independent of the initial tissue size and density, especially considering that cell density evolution shows opposite trends at early stages of expansion for small and large tissues (Fig. 1C). This observation suggests that the early stages of epithelial expansion are primarily driven by cell migration rather than proliferation or density-dependent decompression and cell spreading.

The observation that *υ_r_* is independent of tissue size ought to explain why small tissues have faster area expansions than large tissues. We hypothesized that the relation between tissue size and areal increase could be attributed primarily to the perimeter-to-area ratio. Assuming a constant edge velocity *υ_n_* normal to the tissue boundary, the tissue area increases as *dA* = *Pυ_n_dt*, where *P* is the perimeter of tissue and *dt* is a small time interval. Thus, the relative area increase *dA/A* = (*P/A*)*υ_n_dt* scales as the perimeter-to-area ratio, which is inversely proportional to the radius for circular tissues, so the relative area increases faster for smaller tissues (Fig. 1B). In order to verify that the perimeter-to-area ratio is proportional to the relative area increase, we analyzed elliptical tissues with the same area and tissue density but different perimeters (Movie S3). Increasing the perimeter-to-area ratio of a tissue by increasing its aspect ratio indeed leads to faster areal expansion (Fig. 1E). A simple, edge-driven expansion model with linear increase of the tissue major and minor axes predicts *A*(*t*)/*A*(0) = (*a* + *υ_n_t*)(*b* + *υ_n_t*)/(*ab*), where *a* and *b* are the initial major and minor axes of the tissue. This model fits our data well assuming the same edge speed *υ_n_* ≃ 29.5 *μ*m/h for all tissues (Fig. 1E). Together, our findings demonstrate that epithelial shape and size determine area expansion dynamics via the perimeter-to-area ratio. This relationship results from the fact that tissues exhibit a constant, size-independent, migration-driven edge speed normal to tissue boundary. Since initial tissue size does not affect boundary dynamics, but does impact the relative growth and expansion of the tissue, we hypothesize that cells in the tissue bulk exhibit tissue size-dependent behaviors.

### Spatiotemporal dynamics of migration speed and radial velocity

Having demonstrated the role of the boundary in the expansion of large-scale epithelia, we sought to relate tissue areal expansion rate to internal collective cell migration dynamics. We used Particle-Image-Velocimetry (PIV, See SI Appendix) to obtain flow fields describing cell migration within freely expanding epithelia (6, 17, 27, 38, 39). We constructed kymographs (SI Supplementary Methods) to display the full spatiotemporal flow patterns of the tissue (Fig. 2A, B) (20, 21), averaging over the angular direction and over 16 tissues (for representative kymographs, see SI Appendix, Fig. S2). We also separately show time evolution (Fig. 2C) and spatial profiles (Fig. 2D) of speed and radial velocity to compare small and large tissues.

**Fig. 2.**
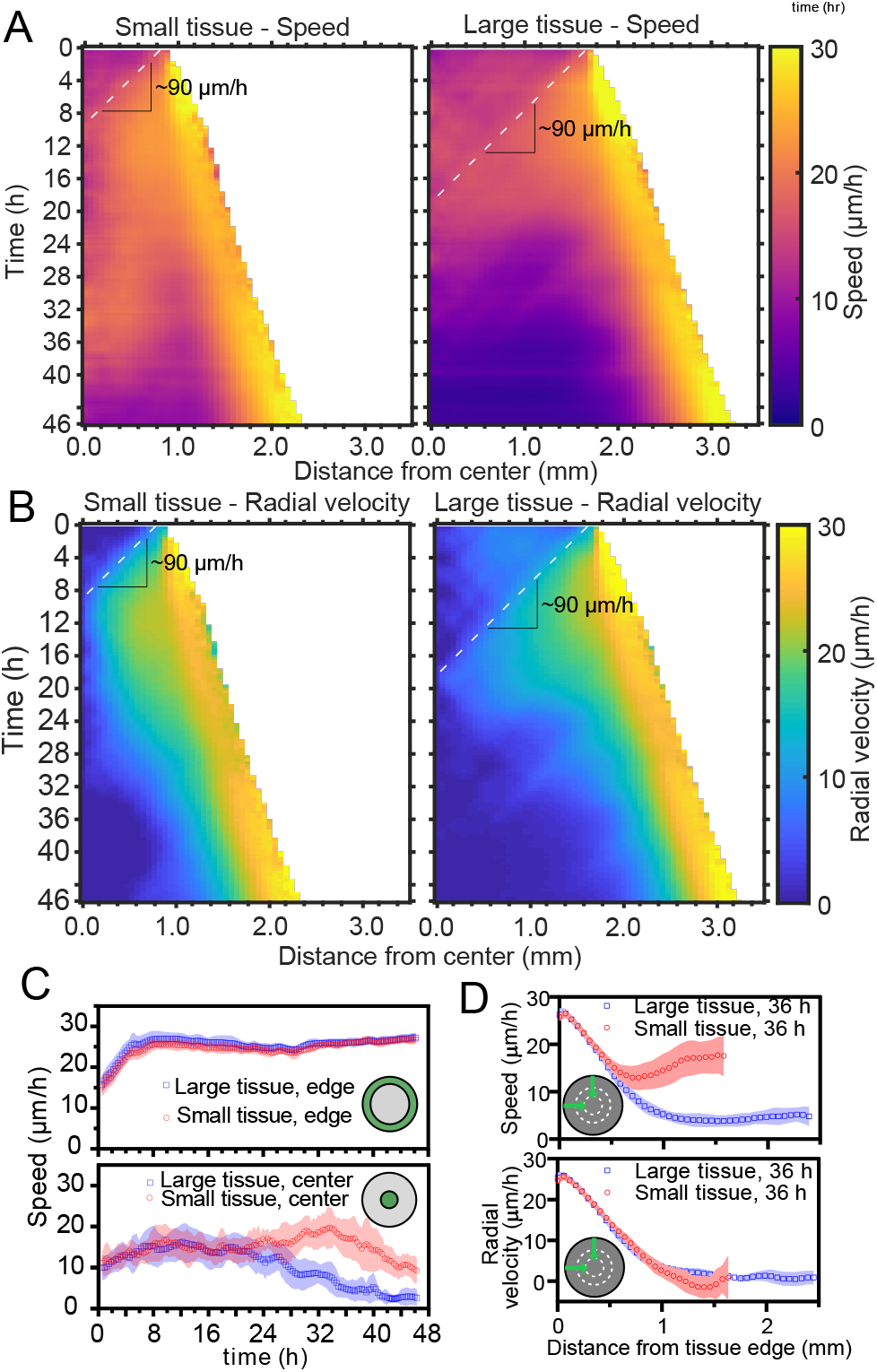
Speed and radial velocity in inner and outer tissue zones. (A,B) Average kymographs of (A) speed and (B) radial velocity *υ_r_* throughout expansion for small (left) and large (right) tissues. (C) Evolution of the average speed of center (top) and boundary (bottom) zones, defined as regions extending ~200 *μ*m from the tissue center and tissue edge, respectively. The width of the zones corresponds roughly to the velocity-velocity correlation length for MDCK cells (17). While the speed in the edge zone remains high in both small and large tissues, the speed in the center zone begins to decrease ~24 h sooner in large tissues than in small tissues, as the central zone of the small tissues has particularly high speed from 18-36 h. (D) Profiles of speed (top) and radial velocity (bottom) at 36 h, from the edge of the tissue inwards. Arrows indicate that the tissues are indexed from the edge of the tissue inwards.

Kymographs of speed and radial velocity reveal the existence of an edge region of fast, outward, radial cell motion (Fig. 2 A, B), with speeds similar to the radial edge velocity reported in Fig. 1D. Up to ~500 *μ*m from the tissue edge, the speed and radial velocity profiles are practically identical for small and large tissues (Fig. 2D), showing that cell motion near the tissue edge is independent of tissue size.

The tissue centers, in contrast, exhibit size-dependent behaviors. For both small and large tissues, a wave front of cell speed and radial velocity propagates toward the tissue centers at ~90 *μ*m/h (Fig. 2A and B, dashed lines). This is approximately 3X faster than the tissue edge speed, consistent with previously described waves of strain rate in cell monolayers (20). Soon after the wave of radial velocity reaches the center, it retreats, leaving a region of low radial velocity that increases in extent in the center of both small and large tissues (Fig. 2B). This decrease of radial velocity is accompanied by a reduction in cell speed in the center of large tissues but not in small tissues, in which cell speed remains high until 36 h (Fig. 2A, 2C Bottom). We examine the behavior of this highspeed but low-radial-velocity central region of small tissues in the next section.

### Emergence of large-scale vortices

The propagation of low radial velocity out from the center of small tissues coincides with the formation and expansion of a millimeter-scale, persistent vortex (see Fig. 3A, Movie S4 for representative vortex). These large vortices are observed in both small and large tissues (Movie S5), but they only reach tissue-spanning sizes in small tissues.

**Fig. 3.**
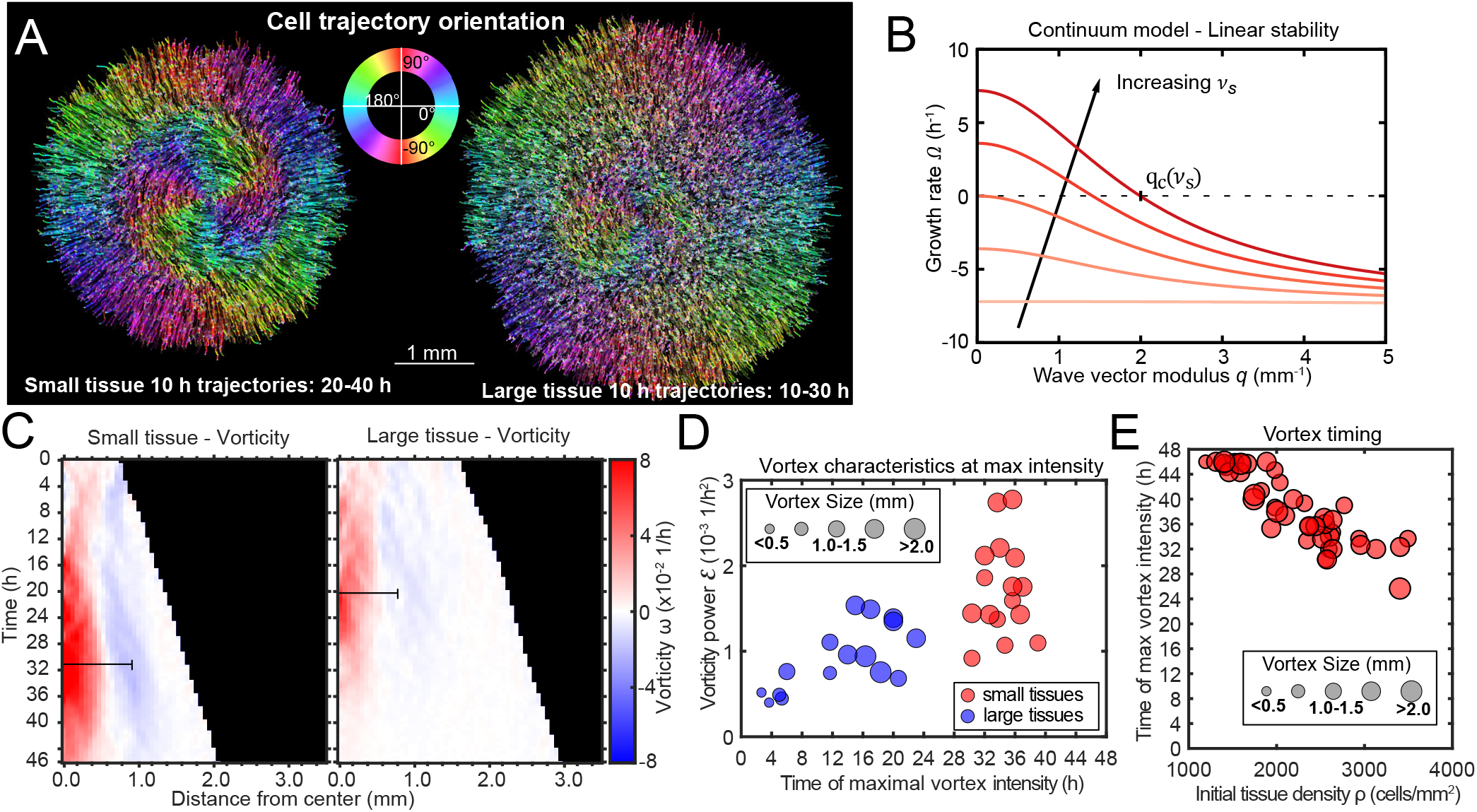
Vortex formation in expanding tissues. (A) Vortical flows seen from 10 h traces of cell trajectories in small (left) and large (right) tissues. We color each trajectory according to its local orientation. (B) Growth rate of perturbations of wave vector modulus *q* around the unpolarized state of the tissue bulk, Eq. 3. Perturbations with wavelength longer than 2*π*/*q_c_* grow (Ω > 0), leading to large-scale spontaneous flows in the tissue bulk. We show curves for the following values of the polarity-velocity coupling parameter: *ν_s_* = 0,1,2,3,4 mm^−1^. For the remaining parameters, we took *T_a_* = 100 Pa/*μ*m, *ξ* = 100 Paos/*μ*m^2^, *η* = 25 MPaos, *γ* = 10 kPaos, *a* = 20 Pa, *K* = 10 nN, as estimated in Ref. (12). (C) Average kymographs of vorticity show that the vortex in small tissues appears in the center and expands to > 1 mm (n = 16), while vorticity in large tissues is generally weaker except during the early stages of tissue expansion (n = 16). The black bars indicate a characteristic vortex size. (D) Characteristic size (marker size), time (horizontal axis), and intensity (vertical axis) of each tissue’s maximal vortex intensity. (E) For small tissues, the time of maximal vortex intensity correlates negatively with the initial cell density.

To visualize the form and scale of these vortices, we tracked individual cell motion and colored cell trajectories according to their orientation (40) for a representative small and large tissue vortex (see Fig. 3A and SI Supplementary Methods). We plotted trajectories for the time periods that the vortex was most apparent, which was 20-40 h in the small tissue (Fig. 3A, left) and 10-30 h in the large tissue (Fig. 3A, right). During the vortex period in small tissues, cell trajectories are primarily radial in the boundary zone, but mainly tangential in the entire central zone (Fig. 3A left, see SI Appendix, Fig. S3 for vortex trajectory quantification).

To understand the emergence of the vortices, we build on a continuum physical model of tissue spreading that describes the cell monolayer as a two-dimensional compressible active polar fluid (12, 37, 41). Consistent with our velocity measurements (Fig. 2C), we assume that cells at the edge zone are radially polarized and motile, whereas cells in the bulk of the tissue are unpolarized and non-motile. We describe cell polarization at a coarse-grained level via a polarity field **p** that obeys the following dynamics (4):

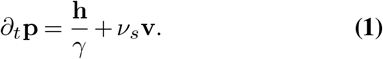

Here, *γ* is the rotational viscosity that damps polarity changes. Respectively, **h** = −*a***p** + *K*▽^2^**p** is the so-called molecular field that governs polarity relaxation: the first term drives the polarity to zero, and the second term opposes spatial variation of the polarity field. As a result of these terms, the radial polarity at the tissue edge decays over a length scale 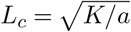 into the tissue bulk.

With respect to previous models of tissue spreading, we add the last term in Eq. 1, which couples the polarity to the tissue velocity field **v**. This coupling is a generic property of active polar fluids interacting with a substrate (42–45). Previous works in agent-based models showed that similar polarity-velocity alignment interactions (4) can lead to waves (14), flocking transitions (46–50), and vortical flows (51–55) in small, confined, and polarized tissues. Here, using a continuum model, we propose that cell polarity not only aligns with but is also generated by tissue flow, and we ask whether this polarity-velocity coupling can lead to large-scale spontaneous flows in the unpolarized bulk of unconfined tissues.

To determine the flow field **v**, we impose a balance between internal viscous stresses in the tissue, with viscosity *η*, and external cell-substrate forces, including viscous friction with coefficient *ξ*, active traction forces with coefficient *T_a_*, and the cell-substrate forces associated with the polarity-velocity coupling *ν_s_*:

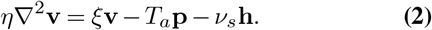

This force balance predicts that even if cell polarity, and hence active traction forces, are localized to a narrow boundary layer of width *L_c_* ~ 50 *μ*m (12, 41), cell flow can penetrate a length 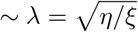 into the tissue. Based on our measurements (Fig. 2D), we estimate λ ~ 0.5 − 1 mm.

A linear stability analysis of Eqs. 1 and 2 shows that perturbations of wave number *q* around the quiescent (**v** = 0) and unpolarized (**p** = 0) state grow with a rate

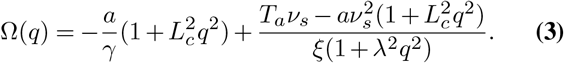

This result shows that, if 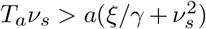, the unpolarized state of an active polar fluid described by Eqs. 1 and 2 is unstable (Ω > 0) to perturbations of wavelength longer than a critical value 2*π*/*q_c_* given by Ω(*q_c_*) = 0 (Fig. 3B). This analysis suggests that, for tissues larger than this critical value ~ 2*π*/*q_c_*, the quiescent tissue bulk becomes unstable and starts to flow spontaneously at large scales, consistent with the emergence of large-scale vortices. The mechanism of this instability is the positive feedback between flow-induced cell polarization and the flows due to migration of polarized cells. The fact that a critical size of the order of millimeters is required for this long-wavelength instability might explain why large-scale vortices have not been observed in previous studies, which considered smaller tissues.

### Vortex kinematics

To quantify the kinematics of the large-scale vortical flows, we obtained the vorticity field *ω* (**r, t**) = ▽ × **v**(**r, t**). Before averaging over tissues, we took the dominant direction of rotation of each tissue to correspond to positive vorticity. This direction was counterclockwise in 51.5% of tissues and clockwise in 49.5% of tissues, with a sample size of 68. With this convention, the vortex core always has positive vorticity. Accordingly, the outer region of the vortex exhibits negative vorticity (Fig. 3C), which corresponds to the counter-rotation that occurs when the central vortical flow transitions to the outer radial flow (Fig. 3A, left). We define a characteristic vortex radius as the radial position of the center of the negative-vorticity region, which is ~1 mm at 36 h in small tissues (Fig. 3C, black bars).

To analyze vortex dynamics across different tissues with varying vortex positioning, and to quantitatively capture the onset and strength of vortices, we calculated the enstrophy spectrum 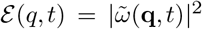, where 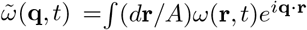 are the spatial Fourier components of the vorticity field *ω*(**r**,*t*) (56). The enstrophy spectrum is the power spectral density of the vorticity field as a function of the wave-vector modulus *q*, and therefore provides a measure of the vortex intensity at a length scale 2*π*/*q*. The kymographs of the enstrophy spectrum show that most of the vortex’s intensity is found at a characteristic length scale of ~1 mm (SI Appendix, Fig. S4).

For each tissue we characterized the maximal vortex strength by the maximum value of 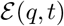 as well as its associated wavelength 2*π*/*q* and time of occurrence. We represented these three quantities on a scatter plot, which shows that vortices in small tissues have generally higher intensity than those in large tissues (Fig. 3D). Vortices in small tissues are also larger relative to tissue size, since the absolute size of vortices in small and large tissues is similar (Fig. 3D). Furthermore, vortex strength peaks several hours later in small tissues than in large tissues (Fig. 3D). We hypothesized that this difference is due to large tissues featuring a faster density increase than small tissues (Fig. 1C). To test this hypothesis, we varied the initial cell density of small tissues and observed that the initial cell density correlates negatively with the time of maximal vortex strength (Fig. 3E, SI Appendix, Fig. S4). These results prompted us to examine spatiotemporal cell density evolution.

### Spatiotemporal dynamics of cell density

Given that cell density appears to affect vortex formation and is known to control contact inhibition of locomotion and proliferation (57), we explored the spatiotemporal evolution of cell density. Constructing average kymographs in the same way as for speed, radial velocity, and vorticity, we observe that the vortex region in the center of small tissues is accompanied by an unexpected local density minimum (Fig. 4A). Strikingly, snapshots of small and large tissues reveal that large-scale vortices nucleate in low-density regions, regardless of location within the tissue (SI Appendix, Fig. S5). However, given that vortices in large tissues are often off-centered, the low-density region does not appear in their average kymograph of cell density (Fig. 4A).

**Fig. 4.**
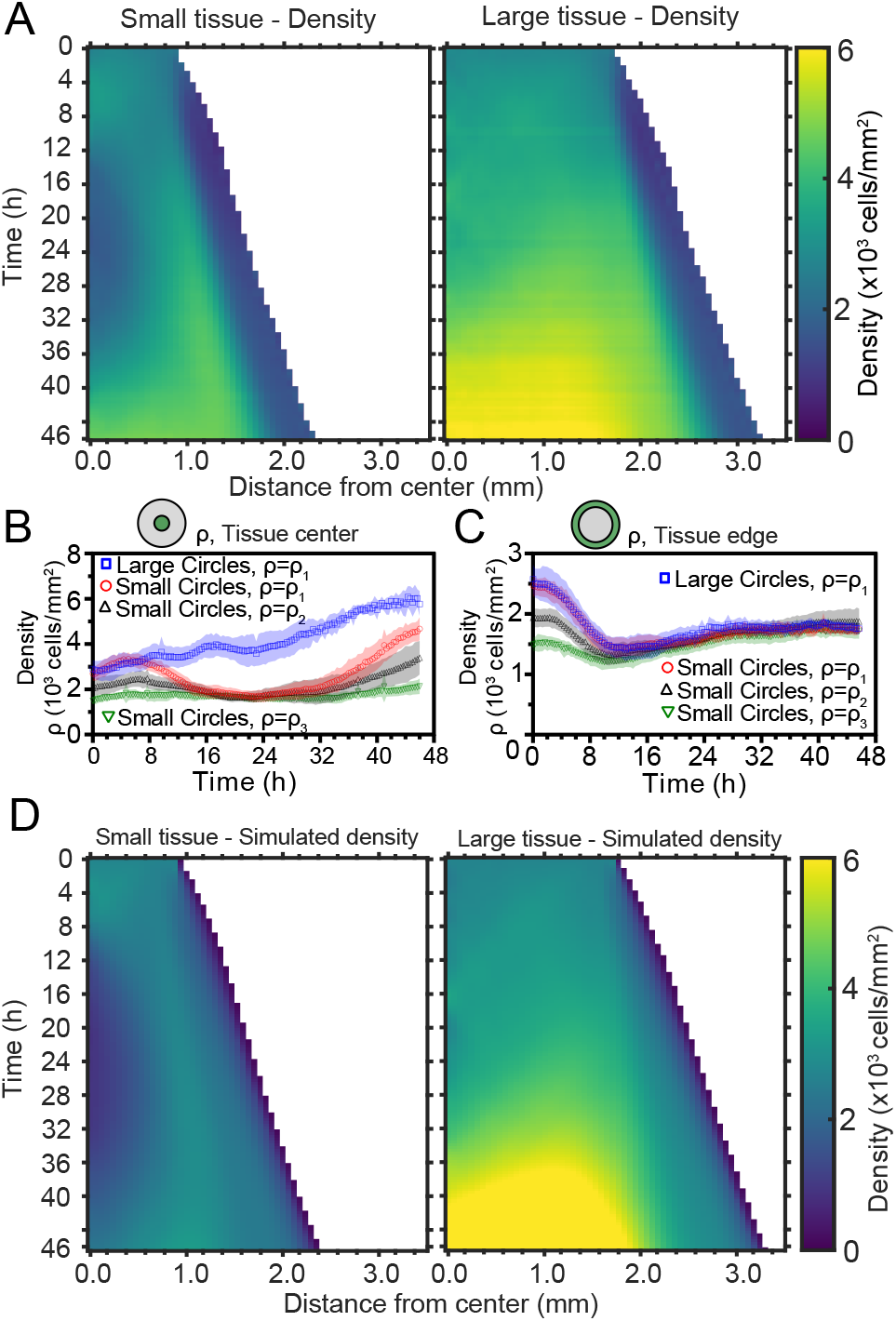
Spatiotemporal dynamics of cell density during epithelial expansion (A) Averaged kymographs of cell density for small (left, n=11) and large (right, n=9) tissues. Small tissues develop a central low-density region that persist more than 20 h. (B) Cell density, *ρ*, at the center of large tissues increases gradually, while cell density at the center of small tissues has non-monotonic evolution. (C) For different initial tissue sizes and densities, the evolution of the cell density, *ρ*, at the boundary zone converges to similar values at about 12 h, which coincides with the end of the overshoot of edge radial velocity in Fig. 1D. Center and boundary zones are defined as in Fig. 2B. (D) Simulated evolution of cell densities obtained from the numerical solution of the continuity equation using the measured radial velocity *υ_r_*(*r,t*) and a uniform and constant cell proliferation rate corresponding to a 16 h cell doubling time. In (B,C) the initial cell density ranges *ρ*_1_, *ρ*_2_, and *ρ*_3_ are the same as in Fig. 1D.

To investigate the effects of initial conditions, we tracked the density evolution of the center and boundary zones across tissues with different starting densities and sizes, grouping initial densities into 3 ranges as before (Fig. 4B and C). As with the average density in Fig. 1C, the density monotonically increases in large tissues centers but is non-monotonic in small tissues. Notably, the cell density at the center of small tissues of different initial cell densities reach a common minimum during the 16-32 h time period (Fig. 4B), which includes the vortex onset time. At the boundary zone, the long-time evolution of the cell density is independent of initial tissue size and density (Fig. 4C). This common long-time evolution is reached at about 12 hours (Fig. 4C), which coincides with the time at which the edge radial velocity stabilizes upon the overshoot (Fig. 1D).

To understand the unexpected transient density decrease at the center of small tissues, we sought to explain it as the result of combined advective transport based on the measured radial flow fields **v**_*r*_(**r**,*t*) and homogeneous cell proliferation at a rate *k*(**r**,*t*) = *k*_0_ throughout the tissue. To test this hypothesis, we solved the continuity equation for the cell density field *ρ*(**r**,*t*),

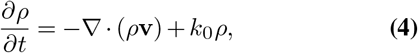

using the average radial velocity profiles *υ_r_*(*r,t*) measured by PIV (Fig. 2D), and a proliferation rate *k*_0_ = 1.04 h^−1^, which corresponds to a cell doubling time of 16 h (SI Appendix, Supplementary Materials and Methods). This minimal model recapitulates the major features of the evolving density profiles for both small and large tissues (compare Fig. 4D with Fig. 4A). Therefore, the unexpected formation of a central low-density region results from the combination of outward tissue flow and proliferation within the colony. However, further research is required to determine the biophysical origin of the non-monotonic density evolution. Moreover, having assumed a density-independent proliferation rate, our model predicts a cell density in the center of large tissues higher than the one measured at the end of the experiment, and it does not quantitatively reproduce the cell density profiles at the edge regions. These discrepancies suggest that more complex cell proliferation behavior is required to fully recapitulate the density dynamics in expanding cell monolayers.

### Spatiotemporal dynamics of cell cycle

To better understand how tissue expansion affects cell proliferation, we analyzed the spatiotemporal dynamics of cell-cycle state. Our cells stably express the FUCCI markers, meaning that cells in the G0-G1-S phase of the cell cycle (referred to here as G1) fluoresce in red (shown as magenta), and cells in the S-G2-M phase of the cell cycle (referred to here as G2) fluoresce in green (34). Additionally, immediately-post-mitotic cells do not fluoresce and appear dark. Small and large tissues are initially well mixed with green and magenta cells, confirming that cells are actively cycling throughout the tissue at the time of stencil removal (SI Appendix, Fig. S6). During tissue expansion, spatiotemporal patterns of cell-cycling behavior emerge (Fig. 5A, Movie S6).

**Fig. 5.**
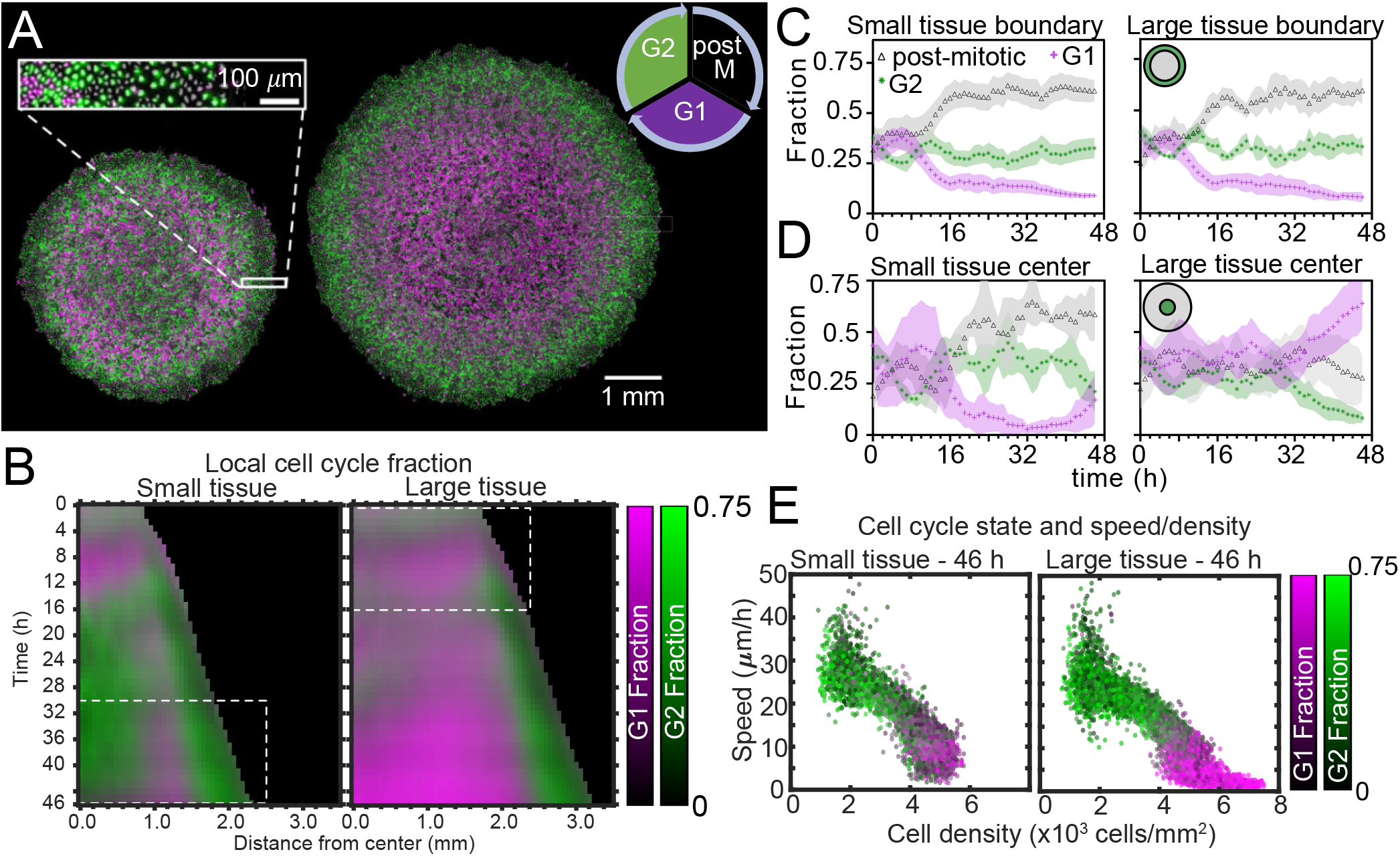
Coordinated spatiotemporal cell-cycle dynamics. Transition from the G1 (magenta) to the G2 (green) phase of the cell cycle corresponds to DNA replication (during S phase). Subsequently, a cell proceeds to mitosis (M phase, dark), and eventually back to the G1 phase upon cell division. (A) Fluorescence images of the Fucci marker of cell-cycle state at the end of the experiment (46 h) of representative small and large tissues overlaid with nuclei positions (gray). The boundary zone of both tissues has more cells in the G2 than in the G1 phase, along with a substantial proportion of dark cells (inset). Scale bars 1mm. (B) Average kymographs (small, n=5; large, n=11) of cell-cycle-state fraction. In small tissues, a G1-dominated transition zone, which appears as a vertical magenta streak from 16 h onward, is interposed between G2-dominated center and edge zones. While the size of small tissues from 30 to 46 h matches that of large tissues from 0 to 16 h (dashed boxes), cell-cycle states between these times are clearly distinct. (C) Fraction of cell-cycle states in the boundary zone. (D) Fraction of cell-cycle states in the center zone. Center and boundary zones are defined as in Fig. 2. (E) Scatter plot of density and speed, with color indicating the fraction of cells at G1 and G2, corresponding to each PIV pixel of the final timepoint of a representative small (left) and large (right) tissue.

To quantitatively investigate these cell-cycle patterns, we obtained the local fractions of G1, G2, and post-mitotic cells by evaluating cell cycle state for each cell nucleus (SI Materials and Methods). We then overlaid kymographs of the G1 and G2 cell-cycle-state fractions (Fig. 5B) and plotted the time evolution of G1, G2, and post-mitotic fractions together (Fig. 5C, D). Immediately after stencil removal, we observe a cell division pulse in all tissues, which manifests in a decrease in G2 and increase in post-mitotic fraction (Fig. 5C,D). After about 12 h of tissue expansion, the boundary region becomes primarily populated by rapidly-cycling cells (Fig. 5B, C), which results in a predominance of cells in this region that either have recently divided (post-mitotic, black) or are likely to divide soon (G2, green). The high numbers of post-mitotic cells indicate that cells in G1 rapidly proceed to mitosis.

In the central region of small tissues (Fig. 5B left, D left), we observe cell-cycling dynamics similar to the boundary region. Thus, in the tissue-spanning vortex of small tissues, cells are also rapidly cycling. The fraction of cells in G1 only starts to increase at ~40 h (Fig. 5D left), coinciding with the weakening of the vortex (Fig. 3C left). In contrast, the center zone of large tissues undergoes strong cell-cycle arrest at the G1-G2 transition at about 30 h, also coinciding with the weakening of the vortex in large tissues (Fig. 5B right, D right). Cells already past G1 at this time continue to division and re-enter G1, evidenced by the steady increase in local fraction of G1 accompanied by a steady decrease in G2 after 30 h. Similar cell-cycle arrests were previously reported both in growing epithelia (35) and in spreading 3D cell aggregates (31). Before the onset of cell-cycle arrest, the center of large tissues exhibits large-scale coordinated cell-cycling dynamics in the form of anti-phase oscillations, with peaks in G2 fraction accompanied by troughs in G1 fraction (Fig. 5B right, D right).

Finally, we sought to link cell-cycle dynamics to the kinematics of tissue expansion by studying correlations between local measurements of cell cycle, cell speed, and cell density (Fig. 5E). Here, each point represents one PIV window, with color indicating its average cell-cycle state. As expected, cell speed is negatively correlated with cell density. Further, in large tissues, the cell-cycle state transitions from G1-dominated to G2-dominated when cell density increases above ~5000 cells/mm^2^ and cell speed falls below ~12 *μ*m/h (Fig. 5E right). In this regime, the decrease of cell speed with increasing cell density bears similarities to previously-reported glass transitions and contact inhibition of locomotion (58–60). Small tissues, by contrast, lack the G1-dominated, slow, high-density cell population (Fig. 5E, left) found in the center of large tissues. Taken together, our findings emphasize that cell cycling, cell flow, and cell density patterns are inextricably linked and depend on the initial size of an expanding tissue.

## Discussion

We began this study by asking how changes in initial size affect the long-term expansion and growth of millimeter-scale epithelia. By means of high spatiotemporal resolution imaging and precisely controlled initial conditions, our assays systematically dissected tissue expansion and growth from the overall boundary kinematics (Fig. 1) to the internal flow patterns (Figs. 2, 3, 4) and cell-cycle dynamics (Fig. 5). While we demonstrated that ‘small’ tissues increase in area relatively much faster than do ‘large’ tissues, our data suggest a surprising and stark decoupling of the outer and inner regions of an expanding epithelium. Notably, the behaviors of the edge zones are largely independent of tissue size, cell density, and history, while interior dynamics depend strongly on these factors.

Unexpectedly, the overall tissue growth and expansion dynamics (Fig. 1) could be attributed to one dominant feature: these epithelia expanded at the same edge speed regardless of initial tissue size, shape, and cell density. As a result of this robust edge motion, the areal expansion rate of the tissue is dictated by its perimeter-to-area ratio. To further emphasize the decoupling of the boundary and internal dynamics of epithelia, consider that the key findings in Fig. 1 neither predict nor depend upon the radically different internal dynamics we observed within ‘small’ and ‘large’ tissues. For instance, despite the roiling vortices occupying large portions of ‘small’ tissues and the pronounced, large-scale contact inhibition of ‘large’ tissues–two antithetical phenomena–no hints of these behaviors can be detected in the motion of the boundary.

Interestingly, the type and timing of internal dynamics are dictated not by the current size but by the expansion history of a given tissue. For instance, while a small tissue eventually expands to reach the initial size of a large tissue, it exhibits different internal dynamics from the large tissue at this size. This difference in internal dynamics is easiest to observe in spatiotemporal evolution of cell cycle (Fig. 5B dashed boxes); the small tissue footprint from 30-46 h closely matches the large tissue footprint from 0-16 h, but the cell cycle distribution during these time periods bears almost no similarities. This of course applies to other important bulk properties of the tissue, as cell cycle is tightly linked to cell speeds and density (Fig. 5E). While edge dynamics are stereotyped and conserved across different sizes, our findings suggest that initial tissue size impacts the bulk dynamics by altering the constraints under which the tissue grows. We expect that tissues with sizes between our two choices would exhibit similar edge dynamics and internal patterns that cross over between our small and large tissues.

The vortices are a particularly striking example of such size- and history-dependent internal patterns. Our active fluid model suggests that the vortices emerge from a dynamical instability of the tissue bulk, which occurs when the tissue reaches a critical size. Thus, whereas the instability itself is a bulk phenomenon independent of the tissue edge, edge-driven expansion allows small tissues to reach the critical size that triggers the instability. In addition, our data suggest a strong correlation between vortex formation and the development of non-monotonic density profiles. Not only small tissues exhibited co-occurrence of vortices with density decreases in the tissue center, but also off-center vortices in large tissues were always preceded by a density decrease at the vortex core (SI Appendix, Fig. S5). These findings should foster the development of more detailed models of tissue vortices to understand their relationship to the cell density field. The pronounced decoupling between boundary and internal dynamics in epithelia confers stability to the overall expansion of the tissue, making it robust to a wide range of internal perturbations. From the perspective of collective behavior, we speculate that such robust boundary dynamics may be beneficial in a tissue whose teleology is to continuously expand from its free edges to sheath organ surfaces. Further, the ability to accurately predict epithelial expansion with a single parameter, the edge speed, will have practical uses in experimental design and tissue-engineering applications. Finally, given that many of the phenomena presented here only occurred due to the millimetric scale of our unconfined tissues and the long duration of the experiments, our results showcase the value of pushing the boundaries of large-scale, longterm studies on freely-expanding tissues.

## Materials and Methods

### Cell culture

All experiments were performed with MDCK-II cells expressing the FUCCI cell-cycle marker system (35). We cultured cells in customized media consisting of low-glucose (1 g/L) DMEM with phenol red (Gibco, USA), 1 g/L sodium bicarbonate, 1% streptomycin/penicillin, and 10% FBS (Atlanta Biological, USA). Cells were maintained at 37°C and 5% CO_2_ in humidified air.

### Tissue patterning

We coated tissue-culture plastic dishes (BD Falcon, USA) with type-IV collagen (MilliporeSigma, USA) by incubating 150 *μ*L of 50 *μ*g/mL collagen on the dish under a glass coverslip for 30 minutes at 37°C, washing 3 times with deionized distilled water (DI), and allowing the dish to air-dry. We then fabricated silicone stencils with cutouts of desired shape and size and transferred the stencils to the collagen coated surface of the dishes. Stencils were cut from 250 *μ*m thick silicone (Bisco HT-6240, Stockwell Elastomers) using a Silhouette Cameo vinyl cutter (Silhouette, USA). We then seeded the individual stencils with cells suspended in media at 1000 cells/mL. Suspended cells were concentrated at ~ 2.25 × 10^6^ cells/mL and pipetted cells into the stencils at the appropriate volume. Care was taken not to disturb the collagen coating with the pipette tip. To allow attachment of cells to the collagen matrix, we incubated the cells in the stencils for 30 minutes in a humidified chamber before flooding the dish with media. We then incubated the cells for an additional 18 hours to allow the cells to form monolayers in the stencils, after which the stencils were removed with tweezers. Imaging began 30 minutes after stencil removal. Media without phenol red was used throughout seeding and imaging to reduce background signal during fluorescence imaging.

### Live-cell Time-lapse Imaging

All imaging was performed with a 4X phase contrast objective on an automated, inverted Nikon Ti2 with environmental control (37°C and humidified 5% CO_2_) using NIS Elements software and a Nikon Qi2 CMOS camera. Phase contrast images were captured every 20 minutes, while RFP/GFP channels were captured every 60 minutes at 25% lamp power (Sola SE, Lumencor, USA) and 500 ms exposure time. No phototoxicity was observed under these conditions for up to 48 hrs. Final images were composited from 4×4 montages of each dish using NIS Elements.

### Tissue edge radial velocity

Tissues were segmented to make binary masks using a custom MATLAB (Mathworks) script. Tissue edge radial velocity was measured from the binary masks within more than 200 discrete sectors of the tissue; the edge radial velocity of all sectors were averaged to arrive at the tissue average edge radial velocity. Radial velocity at each sector was calculated for each timepoint as the rate of change of the average extent of the boundary pixels of the sector, utilized a rolling average of 3 timepoints (1 h) to account for capture phase offsets resulting from capturing phase and fluorescence images at different frequencies. Sectors originated from the center of each tissue at the initial timepoint and were ~ 20*μ*m wide at the edge of the tissue at the starting point.

### Cell counts

The FUCCI system contains a period after M-phase where cells go dark, making FUCCI unreliable for cell counting. Instead, we developed and trained a convolutional neural network to reproduce nuclei from 4X phase contrast images. The output of this neural network was then segmented in ImageJ to determine nuclei footprints and centroids.

### Tissue PIV and density measurements

Tissue velocity vector fields were calculated from 2×2 resized phase contrast image sequences using the free MATLAB package PIVLab (61) with the FFT window deformation algorithm. We used a 1st pass window size of 64×64 pixels and second pass of 32×32 pixels, with 50% pixel overlaps. This resulted in a 115×115 μm window. Local density was also calculated for each PIV window by counting the number of approximate nucleus centroids in that window. Data from PIV was smoothed in time with a moving average of 3 time points centered at each timepoint as before.

### Average kymographs

First, we constructed kymographs for individual tissues using distance from the tissue center as the spatial index for each measurement window corresponding to a kymograph pixel. We did not plot kymograph pixels for which more than 95% of the measurements at that distance were beyond the tissue footprint. We then averaged the individual tissue kymographs, aligning by the centers.

### Trajectory colorization

We first generated a plot of all relevant trajectories (62) colorized randomly in grayscale using a custom MATLAB (Mathworks) script. We then used the Fiji plugin OrientationJ on this plot to colorize the resulting image according to orientation (40).

### Cell density simulation

To test whether the observed spatiotemporal evolution of density could be explained by flow of material and simple proliferation, we solved the continuity equation for a homogenous tissue in a circular geometry with spatiotemporal evolution of radial velocity as measured from 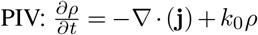, where the current **j** = *ρ***v_r_**.

We discretized the equation in space using the finite volume method, integrating over an annulus with width equivalent to 1 PIV window (~ 115*μ*m). By the divergence theorem, the volume integral becomes a surface integral, and the change in cell number 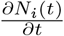 in each annulus is determined by the flux **j** across the inner and outer circular boundaries located at i-1/2 and i+1/2 plus the introduction of cells due to proliferation: 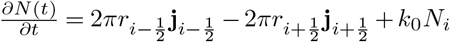.

For numerical stability purposes, we included a diffusion term in the current **j**, using the smallest diffusion coefficient 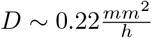 that sufficiently stabilized the density field. 4 The flux was then nothing but the the contribution of current due to radial velocity as well as diffusion as demanded by Fick’s first law: 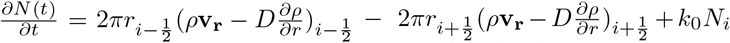.

Dividing by the area of each annulus, we arrive at the discretized equation for the evolution of density for each annulus:

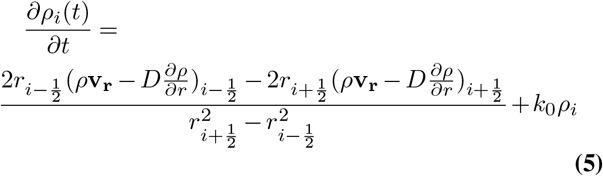

for *i* > 0 and

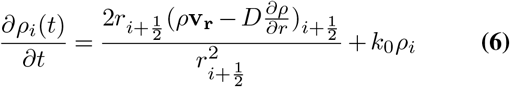

for *i* = 0.

We integrated this equation forward in time with the Euler method using a time step of 20 minutes to align with experimental data collection. We used initial condition 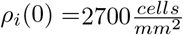 for *r_i_* < *r_tissue_* and *ρ_i_*(0) = 0 for *r_i_* > *r_tissue_*. For comparison with experimental data, we thresholded the kymographs of simulated density at 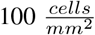, which corresponds to much lower density than a confluent tissue.

### Cell cycle analysis

The Fucci system consists of an RFP and GFP fused to proteins Cdt1 and Geminin, respectively (34). Cdt1 levels are high during G1 and low during the rest of the cell cycle, while Geminin levels are high during the S, G2, and M phases (34, 35). After capturing the appropriate fluorescence images, preprocessing was implemented identically for GFP and RFP channels to normalize channel histograms. To determine local cell cycle fraction, we determined the median value of RFP and GFP signal for each cell nucleus and manually selected thresholds for RFP and GFP signals separately to classify cell cycle for each cell as G0-G1-S (RFP above threshold), S-G2-M (RFP below threshold and GFP above threshold), or postmitotic (RFP and GFP below threshold). Local cell cycle fraction of each state could then be easily computed for each PIV pixel. Note that S phase (both RFP and GFP signals above threshold) did not prove to be a reliable feature for segmentation.

## Acknowledgements

D.J.C. acknowledges the National Institutes of Health R35 GM133574-01. R.A. acknowledges support from the Human Frontiers of Science Program (LT000475/2018-C).

