## Supplemental Information for "Size-dependent patterns of cell proliferation and migration in freely-expanding epithelia"

### Table of contents

**1. Supplementary Figures and Captions** ........ S2

**2. Supplementary Movies** ........ S6

1. Supplementary Figures and Captions

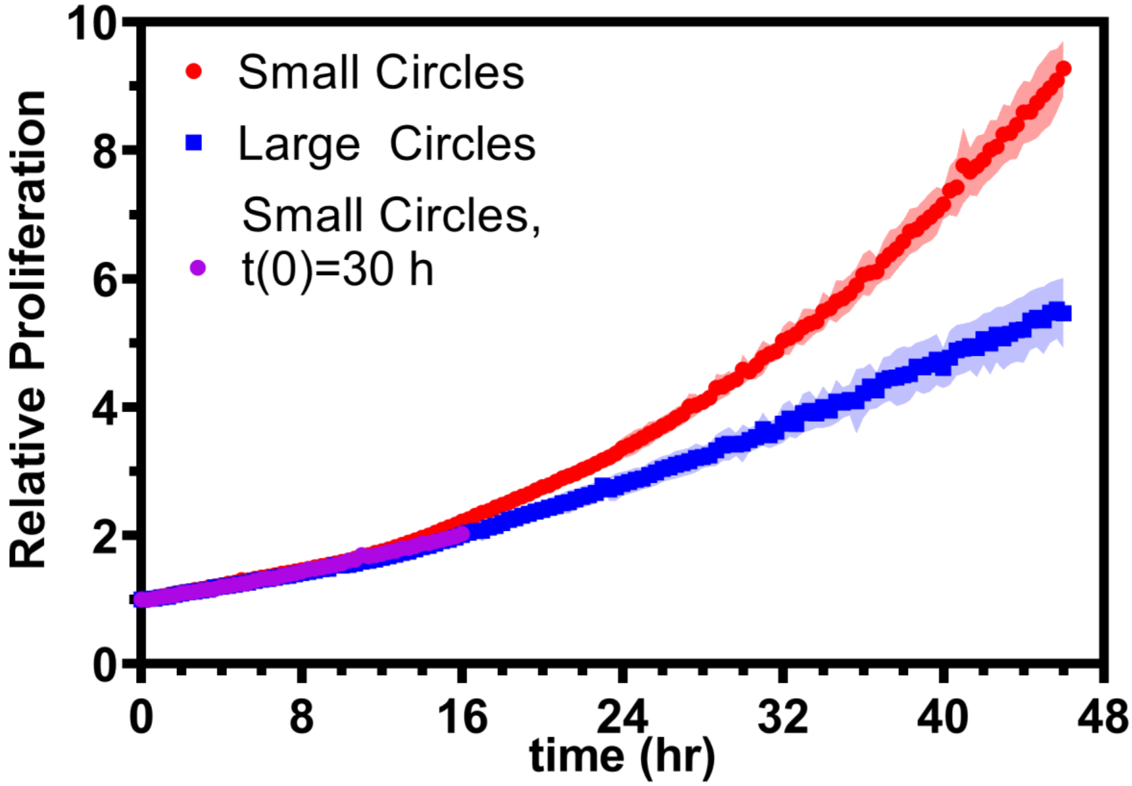


**Figure S1.** Relative proliferation N(t)/N(0) for small and large tissues. Purple points show the relative area increase, $A(t+t_0)/A(t_0)$, of small tissues from the time $t_0$ when they reached the starting size of the large circles.

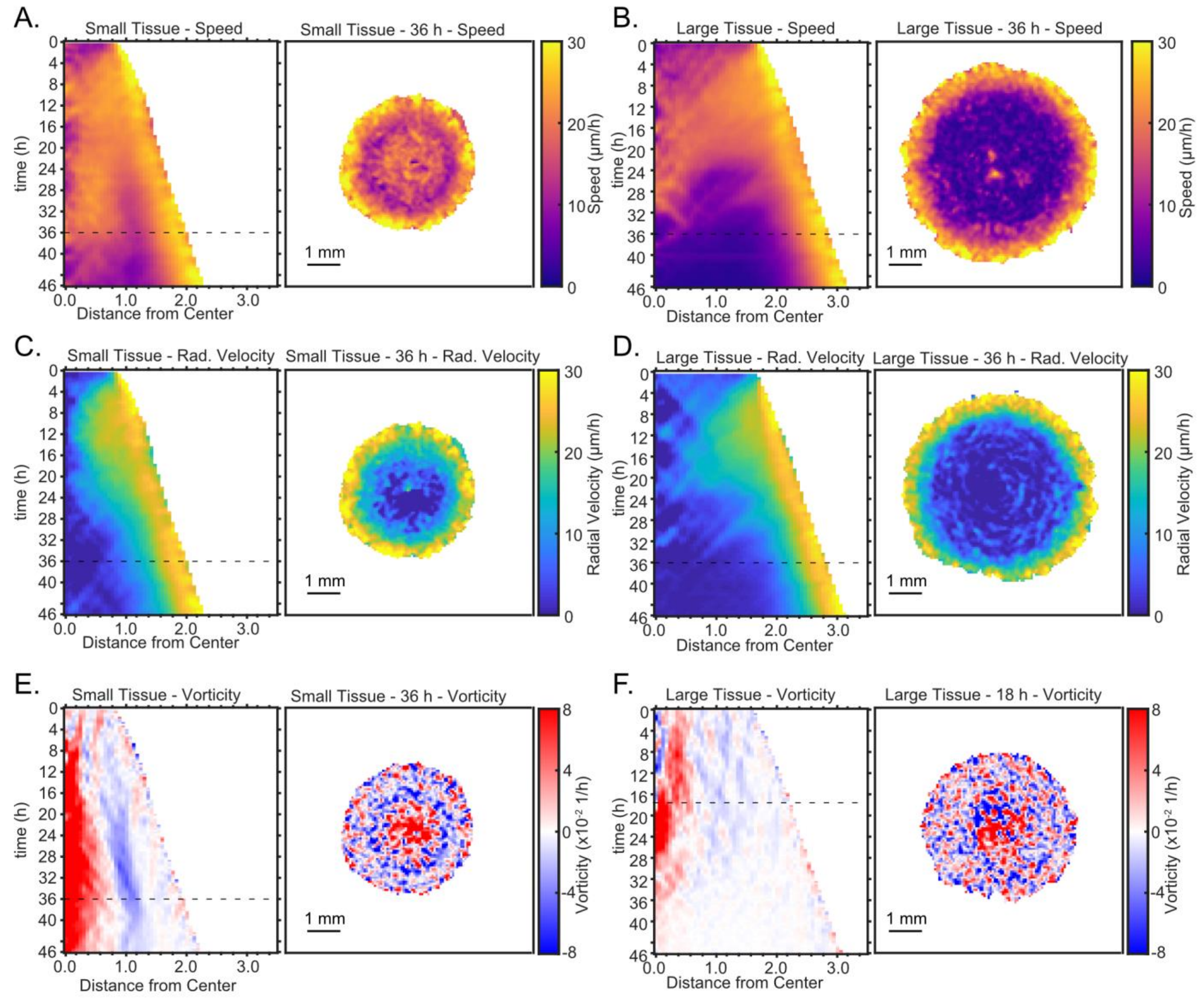


**Figure S2.** Representative kymographs and heatmaps for speed (A-B), radial velocity (C-D), and vorticity (E-F). Dashed lines indicate timepoint to which heatmaps correspond. Kymographs and heatmaps in each column are from the same representative tissue. Note that the snapshot in F is taken from 18 h instead of 36 h to include the time of high vorticity at that earlier timepoint relative to 36 h.

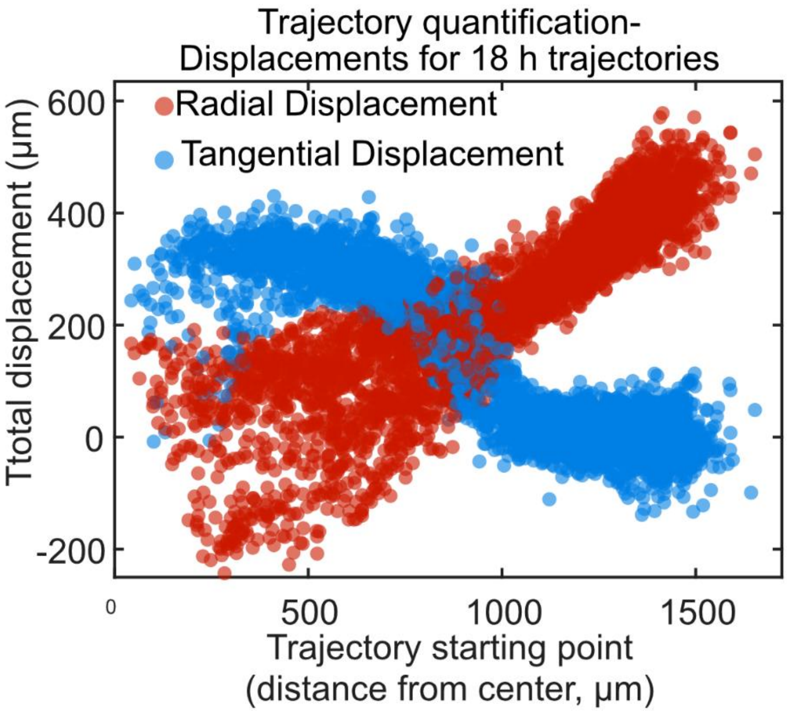


**Figure S3**. Radial and tangential displacements of 18 h cell trajectories for a representative small tissue as a function of the initial radial position of each cell trajectory. Trajectories were selected from the vortex-dominated period of 24-42 h (Supplementary Methods). The tangential displacement (blue markers) most clearly reveals the presence of the vortex, with $\sim 350\mu m$ tangential displacement throughout the central $750\mu m$ of the tissue. At the outer zone, tangential displacements drop to $\sim 0\mu m$. In contrast, radial displacement (red markers) increases roughly linearly from the center to the outer edge of the tissue, with no sharp change when moving from the central to the outer region. The fact that radial displacements are largely insensitive to the vortical flows explains why the presence of a vortex has no noticeable impact on the overall expansion of the tissue. Together, the radial and tangential displacements of the tissue reveal a spiraling vortical flow that combines tangential shear with radial expansion. For cell displacements of the small tissue vortex, we selected the 24-42 h period of trajectories spanning at least that period. We defined radial displacement for each trajectory as:

$$Rad_i = r_{i,t_{42}} - r_{i,t_{24}},$$

Where $r_i$ is the distance from the center of a trajectory, $t_{42}$ is 42 h and $t_{24}$ is 24 h.

We defined angular displacement similarly as:

$$\mathrm{Ang_i} = \theta_{i,t_{20}} - \theta_{i,t_0},$$

Where $\theta_i$ corresponds to the angular location of a cell.

We defined tangential displacement by integrating the tangential component of the trajectory velocity over the course of the trajectory:

$$\tan_i = \int_{t=t_{24}}^{t_{42}} |\vec{v_i}| \cos(\theta_{tan,i})\, dt$$

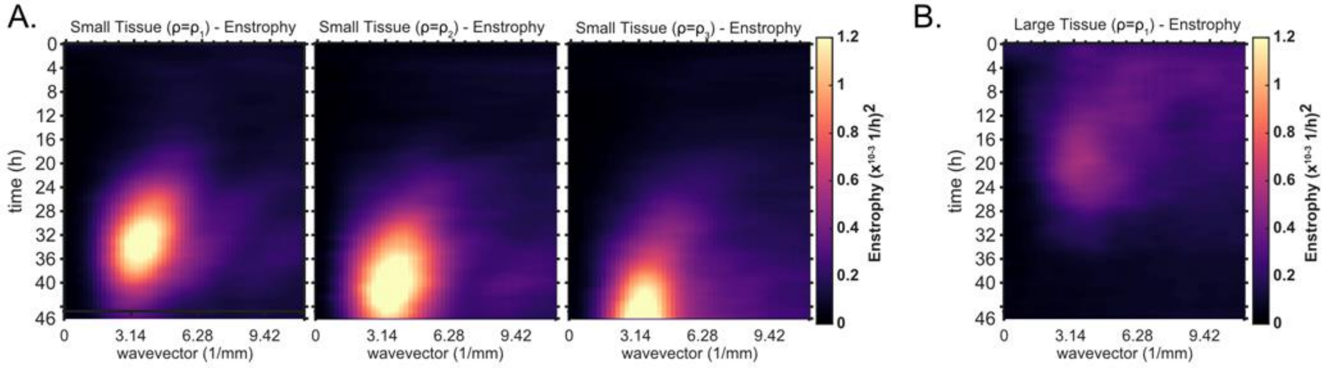


**Figure S4.** Averaged kymographs of enstrophy, which partition the vorticity power in modes of different wavelengths for each timepoint. (A) From left to right, enstrophy kymographs of small tissues for decreasing starting density. $\rho_1 > \rho_2 > \rho_3$. $2350 < \rho_1 < 3050\,\frac{cells}{mm^2}$, $1650 < \rho_2 < 2350\ \frac{cells}{mm^2}$, and $\rho_3 < 1650\ \frac{cells}{mm^2}$,. Decreasing starting density clearly delays the onset of high power, long wavelength (small wavevector) vorticity. Small tissue enstrophy peaks at a wavevector of $\sim 3.14$ mm$^{-1}$, which corresponds to a wavelength of 2 mm. Data from n=16 tissues for $\rho = \rho_1$ (left); n=13 tissues for $\rho = \rho_2$ (middle); and n=11 tissues for $\rho = \rho_3$ (right). (B) Average kymograph of enstrophy for large tissues ($\rho = \rho_1$). The peak at large wavelength is not evident since the vortex is not as prevalent in large tissues.

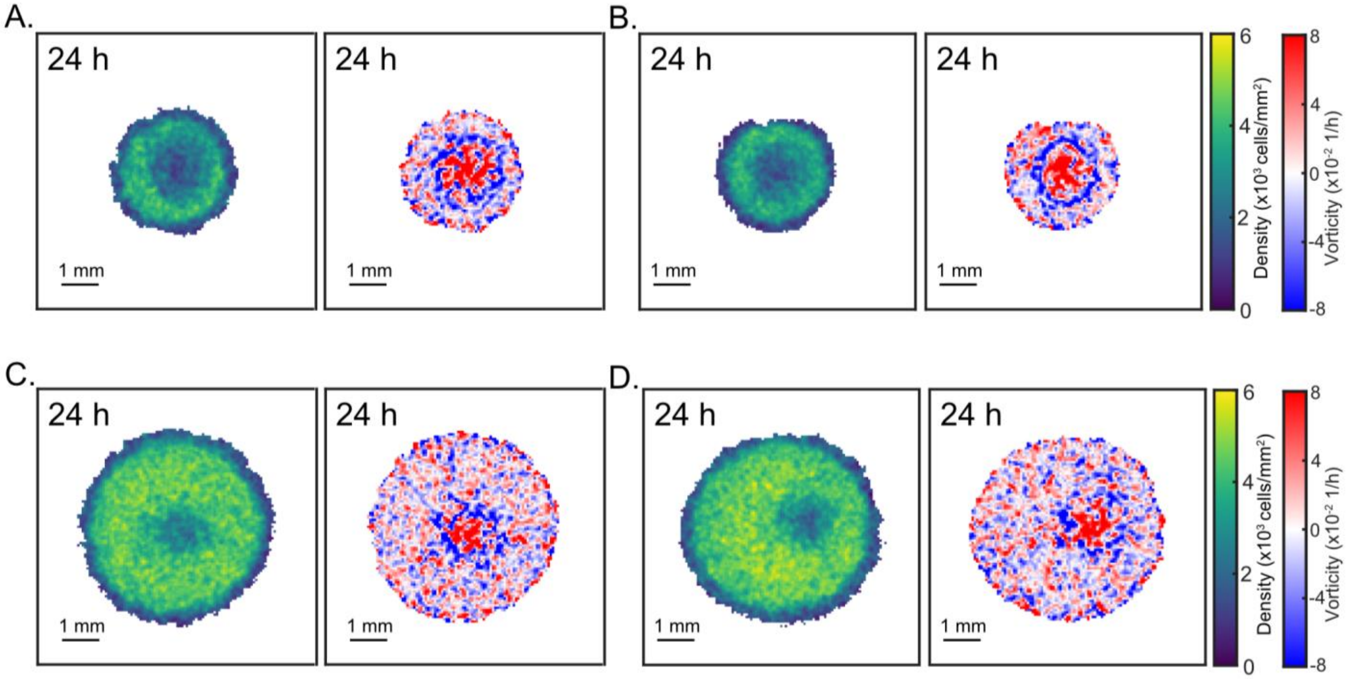


**Figure S5.** Representative heatmaps of density and vorticity for small tissues and large tissues. A region of low density co-occurs with the vortex in representative small and large tissues, centered and off-centered. Representative tissues include small tissues with vortex/low density region in the center-right (A) and center (B) as well large tissues with vortex/low density region the center (C) and right-of-center (D).

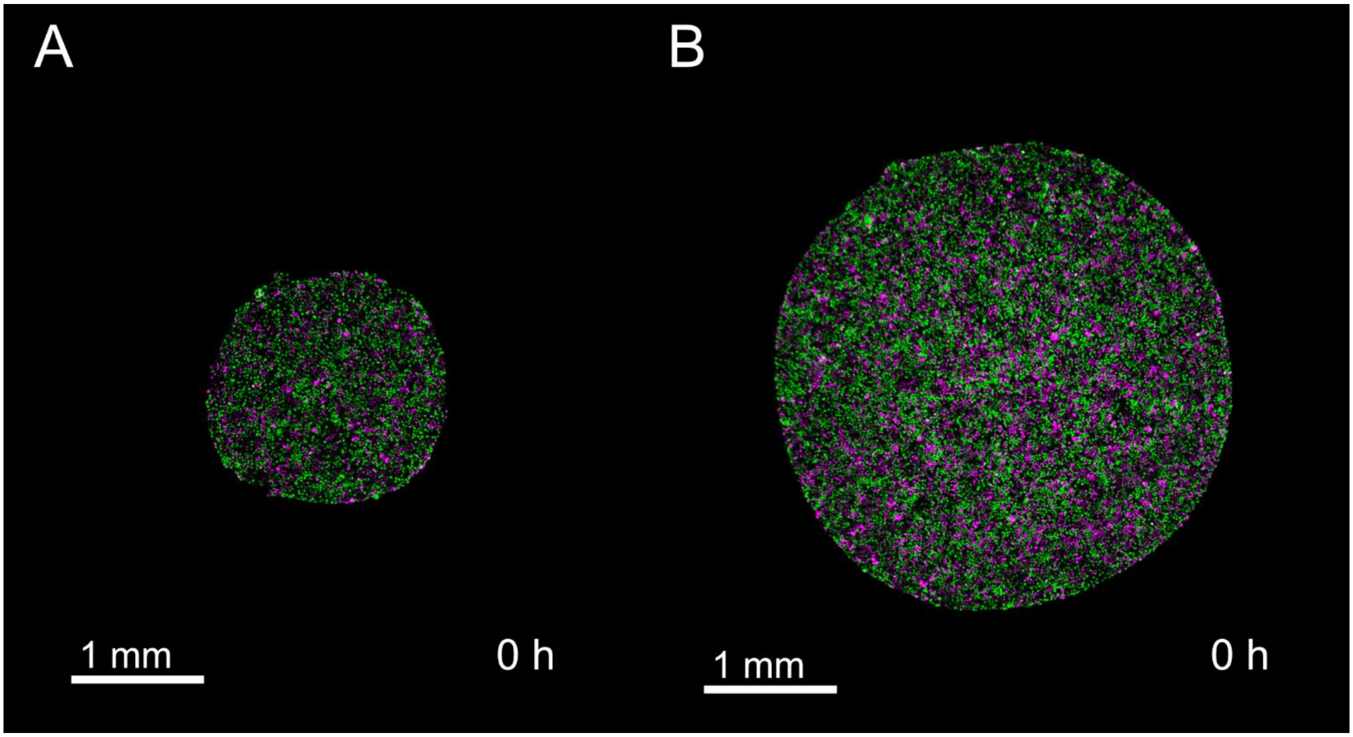


**Figure S6.** Cells within both small and large tissues are actively cycling at the time of stencil removal. (A) Initial timepoint of representative small tissue. (B) Initial timepoint of representative large tissue.

#### 2. Supplementary Movies

**Movie S1.** Effect of size on tissue expansion. Time-lapse phase-contrast microscopy of expansion of representative millimeter-size cell monolayers. The movie shows expansion over a 48 hour period for a sample small (left) and sample large (right) tissues.

**Movie S2.** Time-lapse phase-contrast microscopy showing finger-like protrusions that emerge in the early stages (first 20 hours) of expansion of a cell monolayer.

**Movie S3.** Effect of aspect ratio on tissue expansion. Time-lapse phase-contrast microscopy of expansion of sample elliptical cell monolayers with varying aspect ratios. Area, cell count, and cell density of each elliptical tissue was matched. The major:minor axis ratio of each ellipse was either 1:1, 4:1, or 8:1.

**Movie S4.** Vortex formation in a sample small expanding tissue from t=22 hours to 40 hours of expansion. Left panel shows phase-contrast microscopy of an expanding monolayer. Right panel draws the trajectories of individual cells as they evolve through time.

**Movie S5.** Vortex formation in a sample large expanding tissue from t=12 hours to 30 hours of expansion. Left panel shows phase-contrast microscopy of an expanding monolayer. Right panel draws the trajectories of individual cells as they evolve through time.

**Movie S6.** Coordinated spatiotemporal cell-cycle dynamics in expanding monolayers. Time-lapse fluorescence microscopy of the FUCCI marker for cell-cycle state as the monolayers expand for a small (left) and large (right) tissue over a 48 hour period. Cells in G1 phase are magenta and cells in G2 phase are green.
